# Polyamine transport inhibition and cisplatin synergistically enhance tumor control through oxidative stress in murine head and neck cancer models

**DOI:** 10.1101/2023.07.25.550524

**Authors:** Abdulkader Yassin-Kassab, Nathaniel Wang, Jackson Foley, Tracy Murray Stewart, Mark R. Burns, Robert A. Casero, R. Alex Harbison, Umamaheswar Duvvuri

**Affiliations:** Department of Otolaryngology, University of Pittsburgh School of Medicine, Baltimore, MD, USA; Departments of Oncology and Johns Hopkins University School of Medicine, Baltimore, MD, USA; Departments of Otolaryngology, Sidney Kimmel Comprehensive Cancer Center, Johns Hopkins University School of Medicine, Baltimore, MD, USA; Aminex Therapeutics, Kenmore, WA, USA; The Bloomberg-Kimmel Institute for Cancer Immunotherapy, Johns Hopkins University School of Medicine, Baltimore, MD, USA

**Author notes:** **Data Availability:** The authors confirm that the data supporting the findings of this study are available within the article and its supplementary materials. Correspondence to Umamaheswar Duvvuri, 530 First Avenue, 801.b Smilow Research Center, New York, NY 10016.; R. Alex Harbison, 660 S Euclid, MSC 8115-029-08, Saint Louis, MO 63110. **U. Duvvuri:** 530 First Avenue, 801.b Smilow Research Center, New York, NY 10016. **R.A. Harbison:** 660 S Euclid, MSC 8115-029-08, Saint Louis, MO 63110. **Conflicts of Interest**. M.B. is the president and chief scientific officer of Aminex Therapeutics. R.A.H. and U.D. have a Materials Transfer Agreement with Aminex Therapeutics to receive AMXT-1501 for experimentation. The R.A.C./T.M.S Laboratory has a research contract from Panbela Therapeutics. There are no other conflicts to disclose.

**Keywords:** head and neck cancer, recurrent cancer, cisplatin, polyamines

## Abstract

**Background:** Surgery and/or platinum-based chemoradiation remain standard of care for patients with head and neck squamous cell carcinoma (HNSCC). While these therapies are effective in a subset of patients, a substantial proportion experience recurrence or treatment resistance. As cisplatin mediates cytotoxicity through oxidative stress while polyamines play a role in redox regulation, we posited that combining cisplatin with polyamine transport inhibitor, AMXT-1501, would increase oxidative stress and tumor cell death in HNSCC cells.

**Methods:** Cell proliferation was measured in syngeneic mouse HNSCC cell lines treated with cisplatin ± AMXT-1501. Synergy was determined by administering cisplatin and AMXT-1501 at a ratio of 1:10 to cancer cells *in vitro*. Cancer cells were transferred onto mouse flanks to test the efficacy of treatments *in vivo*. Reactive oxygen species (ROS) were measured. Cellular apoptosis was measured with flow cytometry using Annexin V/PI staining. High-performance liquid chromatography (HPLC) was used to quantify polyamines in cell lines. Cell viability and ROS were measured in the presence of exogenous cationic amino acids.

**Results:** The combination of cisplatin and AMXT-1501 synergize *in vitro* on HNSCC cell lines. *In vivo* combination treatment resulted in tumor growth inhibition greater than either treatment individually. The combination treatment increased ROS production and induced apoptotic cell death. HPLC revealed the synergistic mechanism was independent of intracellular polyamine levels. Supplementation of cationic amino acids partially rescued cancer cell viability and reduced ROS.

**Conclusion:** AMXT-1501 enhances the cytotoxic effects of cisplatin *in vitro* and *in vivo* in aggressive HNSCC cell lines through a polyamine-independent mechanism.

## Introduction

Treatment for head and neck squamous cell carcinoma (HNSCC) is limited to a combination of surgery, radiation, platinum-based chemotherapy, and more recently, immunotherapy for recurrent/metastatic disease. Novel therapies are urgently needed in HNSCC treatment. One such potential therapeutic paradigm involves targeting polyamines, a small family of polycations involved in a multitude of cell functions. Targeting polyamines through inhibition of ODC1, the rate limiting enzyme in polyamine biosynthesis, with difluoromethylornithine (DFMO) in combination with polyamine transport inhibitor, AMXT-1501, has demonstrated promising effects in preclinical studies (1, 2), and is currently undergoing clinical testing in solid tumors (ClinicalTrials.gov Identifier: NCT05500508).

Polyamines are small polycations that regulate cell processes from proliferation and adaptive immunity (3–5) to epigenetic modifications (6, 7), metabolite availability (8, 9), transcriptional regulation (10), and chromatin stabilization (11). PAs also protect the cell from oxidative stress (11–13). Cells both synthesize and acquire PAs from the microenvironment. PAs are immunosuppressive and/or cytotoxic in excess, in part, due to hydrogen peroxide produced by their catabolism (14–18). As tumor cells rapidly turnover, metabolites such as K^+^ (19) and PAs (15) are released which disrupts the extracellular milieu and are immunosuppressive.

Cisplatin chemotherapy is a time-tested mainstay of HNSCC treatment. Cisplatin exerts anticancer activity by forming DNA adducts leading to DNA damage and induces oxidative stress in cancer cells (20, 21) leading to cellular apoptosis (22, 23). Moreover, oxaliplatin downregulates *ODC1* and upregulates *SSAT1* expression, a polyamine catabolic enzyme. This suggests that cisplatin, in part, may exert anti-tumor activity by reducing intracellular polyamines (24). Hypothesizing that the upregulation of polyamine catabolism combined with polyamine transport blockade would diminish tumor cell viability, we sought to test the combination of AMXT-1501 with cisplatin chemotherapy in HNSCC murine models.

## Results

### Cisplatin and AMXT-1501 synergize to enhance cell death *in vitro*

To determine effective *in vitro* concentrations of cisplatin and AMXT-1501, we performed cell proliferation assays in MOC2, TAb2, and MEER PD1R cell lines using a concentration range of 0.08 – 10 µM of CDDP and 2.5 – 80 µM AMXT for 72 hours (Fig. 1A-D). We selected these cell lines as they represent a wide range of aggressive HNSCCs including T cell-depleted, carcinogen-driven (MOC2), genetically altered *TP53* and *PIK3CA* (TAb2), and human papillomavirus-related IZPD-1 resistant (MEER PD1R) head and neck cancer (25–27). A treatment time of 72 hours was used for evaluating changes in cell viability with *in vitro* experiments as MOC2 cells began rapidly proliferating 24 hours after plating (Fig. S1A). MOC2 cells demonstrated increased sensitivity to CDDP compared to TAb2 or MEER PD1R cells (Fig. 1A and 1B). On the other hand, TAb2 cells were more resistant to AMXT than the other cell lines (Fig. 1C and 1D). Next, we tested synergy between combinations of CDDP and AMXT *in vitro*. Using CDDP to AMXT ratios of 1:10, 1:5 and 1:2, Chou-Talalay quantitative analysis of dose-effect relationships was notable for synergy at most concentrations using a ratio of 1:10 (CI < 1.0; Fig. 1E). CDDP and AMXT combination treatment led to synergy for almost all Fa values (proxy for viability) in MOC2 and MEER PD1R cells, while we only observed synergy in TAb2 cells with treatments using concentrations less than 4 µM of CDDP and 40 µM of AMXT. We confirmed synergistic cell death with 2 µM of CDDP and 20 µM of AMXT in the respective cell lines (Fig. 1F-H). We then compared the effect on cell viability of our CDDP and AMXT combination to the efficacy observed in previous studies using AMXT and DFMO (1, 2). When using 3 mM of DFMO, we found that there was no significant difference on cell viability between the CDDP and AMXT combination compared to CDDP and DFMO or when all three drugs were using together (Fig. S2A). As we were interested in identifying a novel treatment combination leveraging cisplatin which is commonly used in the treatment of head and neck cancer, we chose to continue our investigation using the combination of cisplatin and AMXT-1501. Given the finding of synergy between cisplatin and AMXT-1501 in syngeneic murine cell lines *in vitro*, our next goal was to determine if the combination reduced tumor growth *in vivo*.

**Figure 1.**
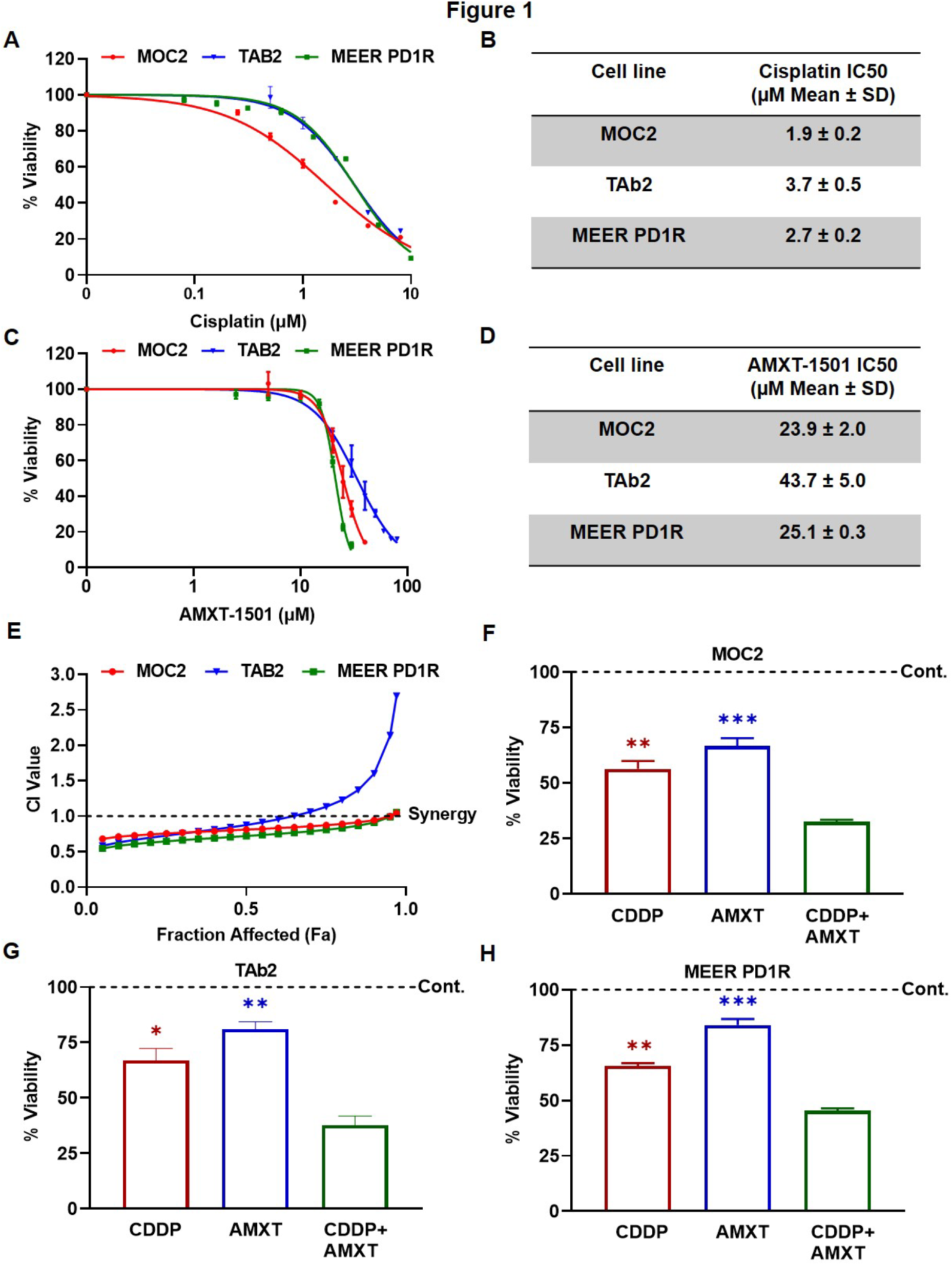
Cisplatin and AMXT-1501 synergize *in vitro*. Cell proliferation assay to determine IC50 values for (A and B) CDDP and (C and D) AMXT in the indicated cell lines after 72h. Representative graph from one experiment is shown. (E) Chou-Talalay dot plot displaying synergy curves with CDDP and AMXT in the indicated cell lines. Representative graph from one experiment is shown. (F-H) Cell proliferation assay in indicated cell lines after treatment with 2 µM of CDDP and 20 µM of AMXT. For F-H, comparisons are made between CDDP + AMXT treatment and individual treatment groups. Statistical significance was calculated using one-way ANOVA with Tukey’s multiple comparison test. *p < 0.05, **p < 0.001, ***p < 0.0001.

### Cisplatin and AMXT-1501 combination *in vivo* reduce tumor growth

To test the effect of CDDP and AMXT alone or in combination on cancer cell proliferation *in vivo*, we transplanted C57BL/6 mice with MOC2 and TAb2 allografts and exposed them to control, CDDP, AMXT, or combination treatment. When using 10 mg/kg/day of AMXT, we found that the combination treatment significantly reduced tumor volume in MOC2 allografts to a greater extent than control or either treatment alone (Fig. 2A and 2B). Similar effects of the combination treatment were observed in TAb2 allografts treated for 7 days (Fig. 2C and 2D). Tumor volume reduction persisted for an additional 6 days after discontinuing treatment in the TAb2 tumors treated with combination therapy.

**Figure 2.**
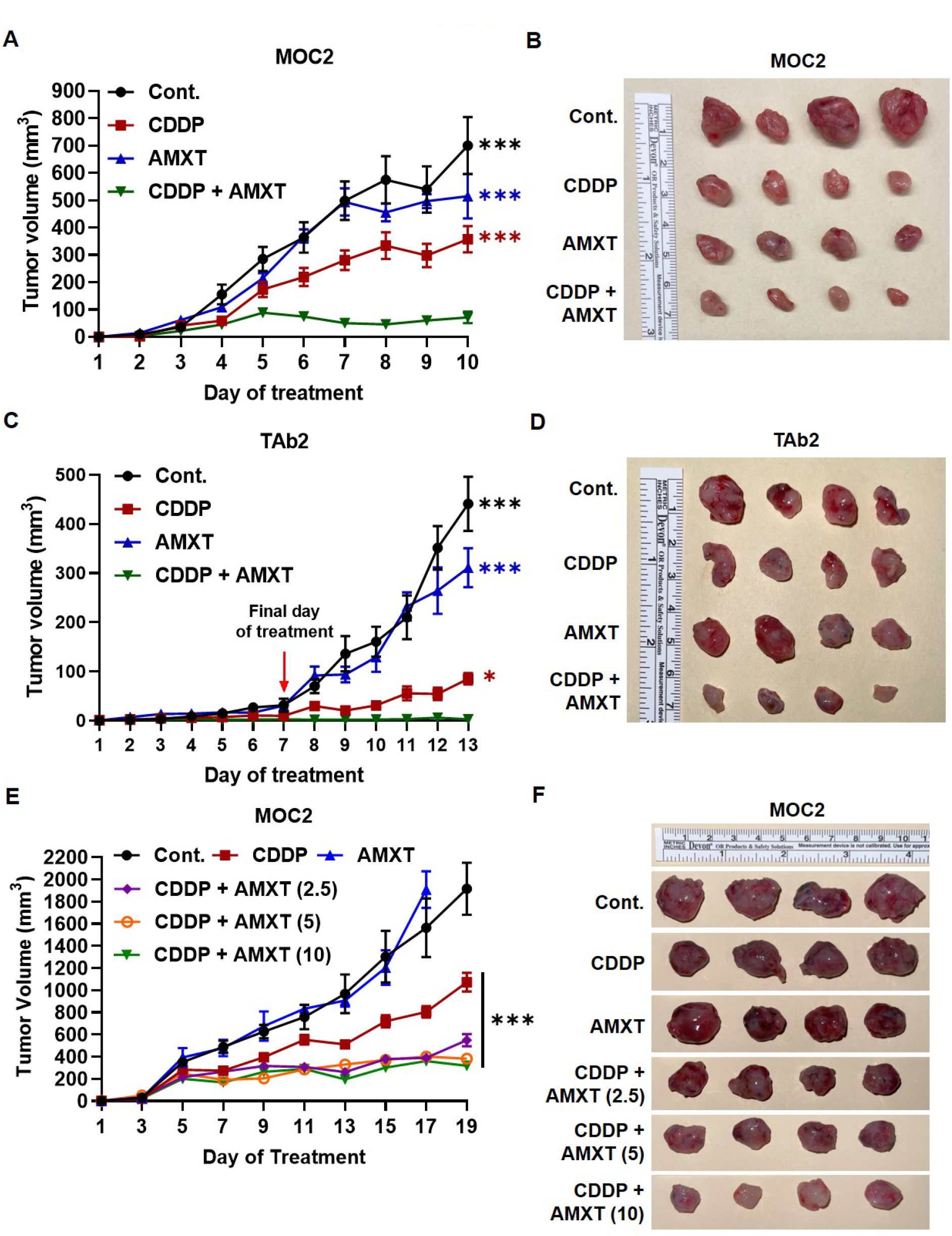
Cisplatin and AMXT-1501 combination synergistically inhibit tumor growth *in vivo*. Tumor growth curves in mice xenografted with (A) MOC2 and (C) TAb2 cells after treatment with CDDP and 10 mg/kg/day of AMXT. Treatment was discontinued in mice harboring TAb2 tumors after day 7. (E) Tumor growth curves in mice xenografted with MOC2 cells after treatment with CDDP and 2.5, 5, and 10 mg/kg/day of AMXT. Mice treated with AMXT alone reached endpoint as defined in the IACUC protocol and were euthanized on day 17. Images for harvested (B and F) MOC2 and (D) TAb2 tumors are shown for each treatment group. For A and C, comparisons are made between CDDP + AMXT treatment and individual treatment groups. For E, comparisons are made between CDDP and all CDDP + AMXT groups. Statistical significance was calculated using two-way ANOVA with Tukey’s multiple comparison test. *p < 0.05, ***p < 0.0001.

Cisplatin is known to have toxic side effects and is specifically known to induce nephrotoxicity, hepatotoxicity, and ototoxicity (28–30). Pathologic analysis was performed on the kidneys of mice treated with cisplatin and AMXT-1501. In the CDDP treatment group, minimal tubular basophilia was observed, a sign of tubular regeneration following acute tubule epithelial injury (31). In the AMXT group (Fig. S3A), there was mild tubular basophilia and minimal tubular necrosis. Moderate tubular basophilia and minimal necrosis was noted in the combination treatment group (Fig. S3B).

To reduce toxicity from treatment, we tested the pharmacodynamic effects of combination treatment using 2.5, 5, or 10 mg/kg/day of AMXT plus four doses of CDDP for 19 days (Fig. 2E and 2F). Pharmacodynamically, we observed significant treatment response in all combination groups, regardless of AMXT dose, compared to control or either treatment alone.

Within the combination groups, we observed a slightly greater response when using 5 or 10 mg/kg/day of AMXT compared to 2.5 mg/kg/day. There was no significant difference in tumor volume observed between the use of 5 or 10 mg/kg/day of AMXT in combination with CDDP. Additionally, we observed no significant difference in varying doses of AMXT with CDDP on body weight compared to CDDP alone.

We also tested the durability of response in MOC2-bearing mice treated with 10 mg/kg/day of AMXT plus four doses of CDDP with treatment discontinuation after 11 days. Similar to the TAb2-bearing mice treated with AMXT plus CDDP, treatment response in the MOC2-bearing mice persisted even after treatment discontinuation (Fig. S4A and S4B). Collectively, these data suggest that CDDP and AMXT act synergistically to diminish tumor proliferation both *in vitro* and *in vivo*.

### Combination of cisplatin and AMXT-1501 induces cell death through apoptosis

Our next objective was to determine the mechanism of cell death caused by CDDP and AMXT combination therapy. Since it has been previously found that both CDDP and AMXT individually induce cell death through apoptosis, we hypothesized that the combination of both drugs would enhance apoptosis (2, 32). Using a TUNEL assay to test for evidence of DNA damage and infer the extent of apoptosis (33) in our harvested tumors, we observed significantly greater TUNEL positivity in the combination treatment group relative to control or AMXT or CDDP alone among the MOC2 (Fig. 3A and 3B) and TAb2 (Fig. S5A and S5B) tumors. To confirm the mechanism of cell death of the CDDP and AMXT combination is through the induction of apoptosis *in vitro*, we utilized Annexin V and propidium iodide (PI) staining. After treatment completion, we found that the percentage of MOC2 cells stained with Annexin V and Annexin V/PI significantly increased in the presence of both drugs (Fig. 3C and 3D). These data indicate that CDDP and AMXT combine to enhance cell death through apoptosis. We next sought to test the hypothesis that AMXT enhances CDDP-induced oxidative stress relative to either agent alone.

**Figure 3.**
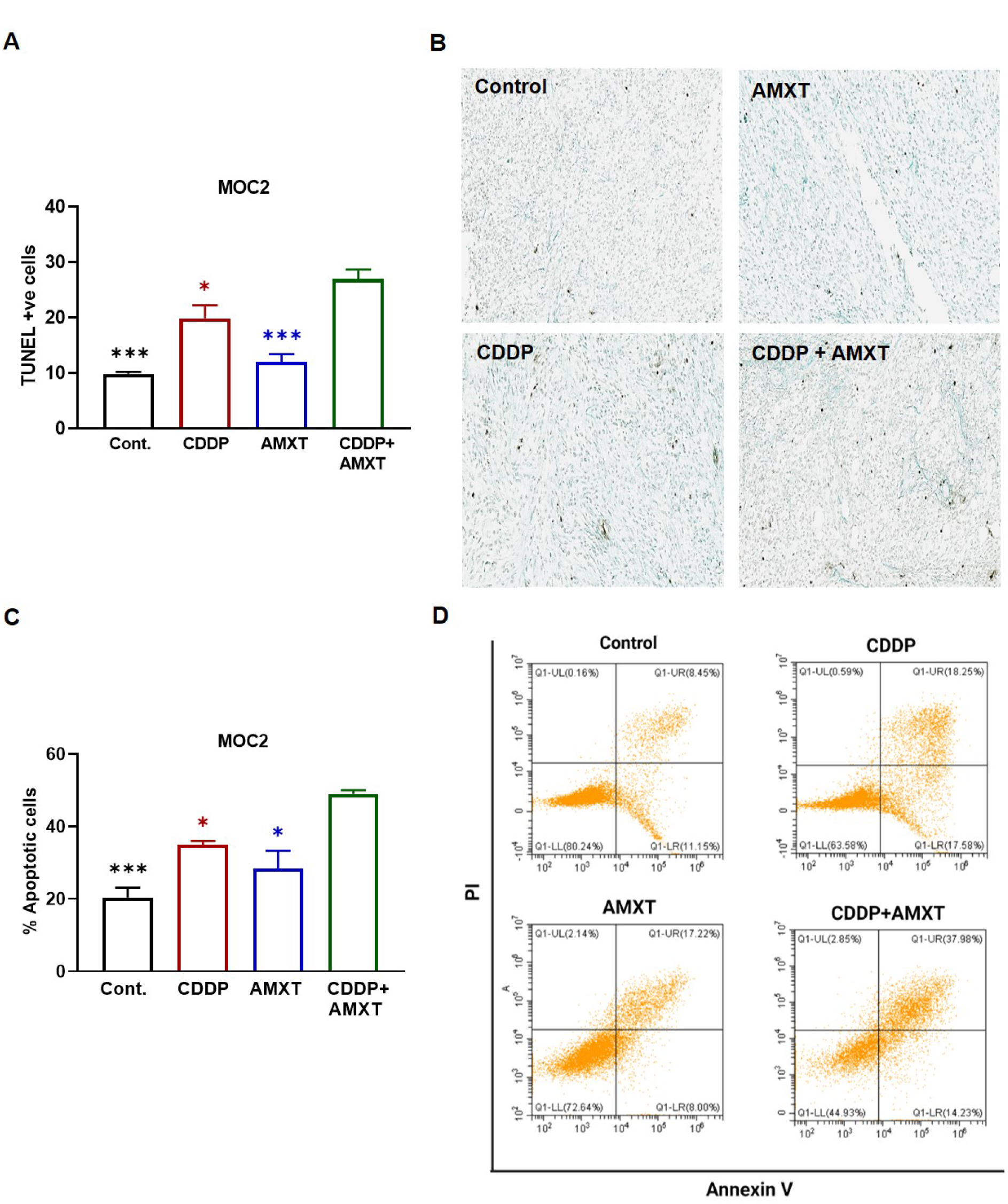
Combination of cisplatin and AMXT-1501 induces cell death through apoptosis. (A) Bar graph analysis of TUNEL staining in harvested MOC2 tumors after treatment with CDDP and AMXT. Bars display the average number of TUNEL-positive cells per high-powered field. (B) Representative TUNEL stain images are shown. (C) Annexin V and PI staining in MOC2 cells after treatment with CDDP and AMXT. Bars display the percentage of cells undergoing apoptosis in each treatment group. (D) Dot histograms from one representative experiment are shown. For A and C, comparisons are made between CDDP + AMXT treatment and individual treatment groups. Statistical significance was calculated using one-way ANOVA with Tukey’s multiple comparison test. *p < 0.05, ***p < 0.0001.

### Combining cisplatin and AMXT-1501 increases reactive oxygen species in MOC2 cells

As polyamines have antioxidant properties (34), we hypothesized that aggressive head and neck cancers, which upregulate oxidative metabolism (35), would exhibit increased susceptibility to the combination of polyamine transport blockade with AMXT-1501 plus cisplatin. To elucidate the mechanism by which CDDP and AMXT synergize, we implemented the DCFDA assay in MOC2 cells and found that reactive oxygen species (ROS) increase nearly 4-fold after combination treatment relative to untreated cells or monotherapy (Fig. 4A). We found that N-acetylcysteine (NAC) abrogates ROS in the combination treatment. We also observed greater production of mitochondrial superoxide in the combination treatment (Fig. 4B).

**Figure 4.**
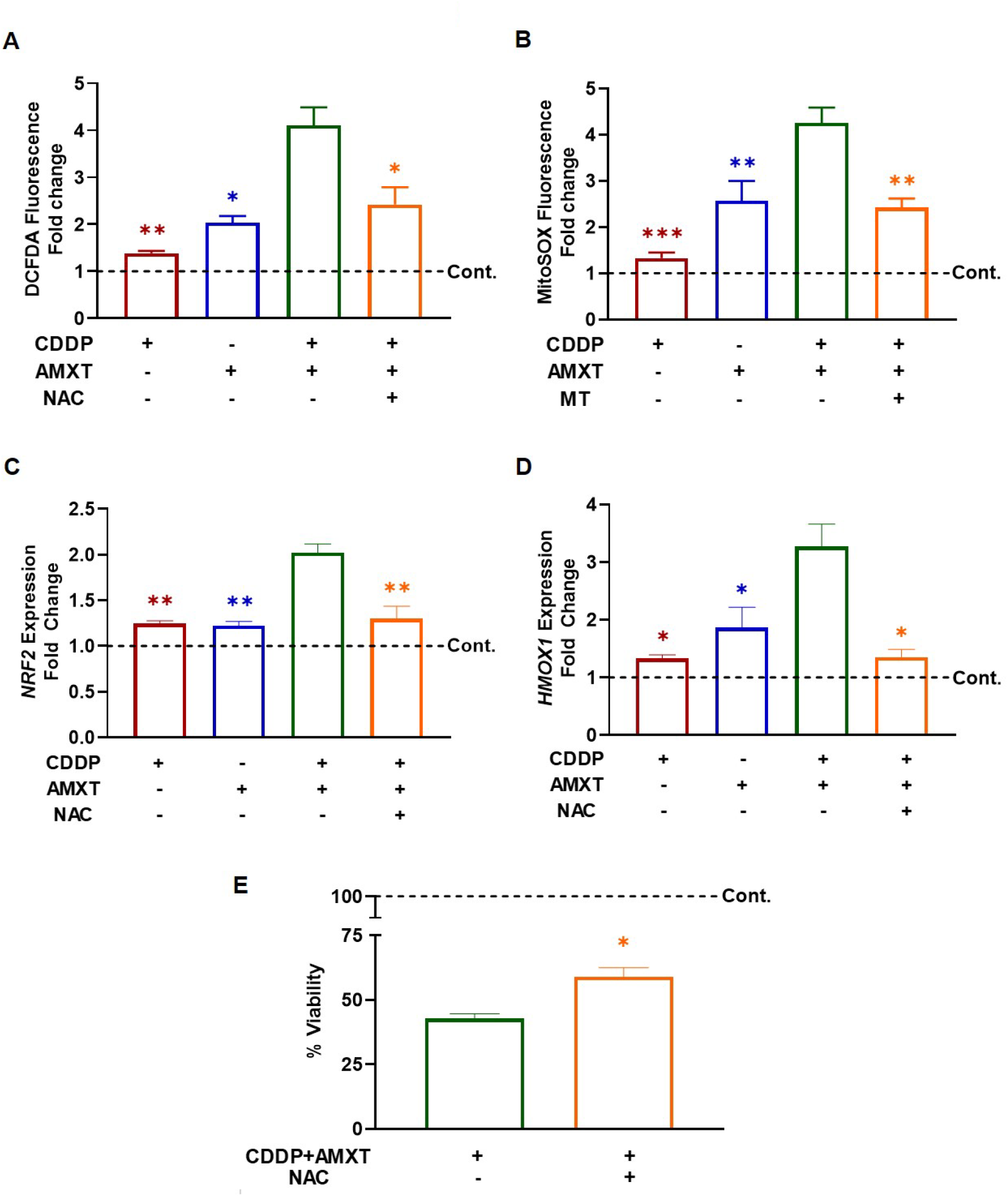
Combining Cisplatin and AMXT-1501 increases reactive oxygen species in MOC2 cells. (A) DCFDA and (B) MitoSOX fluorescence measured in MOC2 cells after treatment with 2 µM CDDP, 20 µM AMXT, 20 mM NAC, and 20 µM of MT for 24h. Expression of (C) *NRF2* and (D) *HMOX1* in MOC2 cells after treatment with CDDP, AMXT, and NAC for 24h. (E) Cell proliferation assay in MOC2 cells after treatment with CDDP and AXMT for 72h with and without the presence of NAC. Comparisons are made between CDDP + AMXT treatment and other treatment groups. Statistical significance was calculated using one-way ANOVA with Tukey’s multiple comparison test. *p < 0.05, **p < 0.001, ***p < 0.0001.

In response to cellular ROS, NRF2 is activated and serves as a transcriptional regulator for other genes involved in protection from oxidative stress, including *HMOX1* (36). To test the effect of ROS on the antioxidant response pathway in the combination treatment, we measured *NRF2* and *HMOX1* expression in MOC2 cells after treatment with CDDP and AMXT for 24 hours. We found *NRF2* expression in MOC2 cells treated with CDDP and AMXT doubled compared to untreated cells (Fig. 4C), while *HMOX1* expression increased by over 3-fold (Fig. 4D), with the expression of both genes significantly higher than their expression after treatment with either drug alone. Additionally, the expression of both genes decreased in the presence of NAC.

To further establish the role of AMXT-1501 in enhancing ROS produced by other drugs known to cause oxidative stress, we performed a DCFDA assay combining AMXT with 10 nM of elesclomol (ES), a potent inducer of oxidative stress (37). The results of our experiment show that combining AMXT and ES significantly increased ROS production compared to either drug alone (Fig. S6A) and that this combination also led to significantly reduced cell viability as well (Fig. S6B).

To test if the increase in ROS observed in the combination treatment is responsible for the synergistic effect on cell death, we repeated the cell proliferation assay with CDDP and AMXT in the presence of NAC and found that viability increased by about 20% (Fig. 4E). Overall, these data suggest that the synergistic effect of cell death of the CDDP and AMXT combination is secondary, in part, to increased cellular ROS.

### Amino acid supplementation diminishes cell death and oxidative stress induced by the combination of cisplatin and AMXT-1501

To determine if the synergistic effect of the CDDP and AMXT combination on cell death and ROS was dependent on changes in intracellular polyamine levels, we performed a rescue experiment by treating MOC2 cells with CDDP and AMXT for 72 hours in the presence of increasing concentrations of spermidine. We found that even with the addition of 1000 µM spermidine, there was no significant change in the viability of the cells after treatment with CDDP and AMXT (Fig. S7A). Treated MOC2 cell pellets were analyzed using HPLC to detect intracellular polyamine levels, however, we found no significant differences in intracellular polyamine content among any of the treatment groups (Fig. S7B-D), indicating that the effects of combination treatment were independent of intracellular polyamine levels.

We then hypothesized that AMXT-1501 may exert cell stress through inhibition of uptake of cationic amino acids given that polyamines are transported, in part, by the cationic amino acid transporter, *SLC7A1* (38). To test this hypothesis, we applied the cationic amino acids: arginine, histidine, and lysine to the media of MOC2 cells treated with CDDP and AMXT. The presence of these amino acids significantly reduced the effects of the CDDP and AMXT combination on ROS (Fig. 5A) and expression of *NRF2* (Fig. 5B) and *HMOX1* (Fig. 5C). In the presence of arginine, histidine, or lysine, MOC2 cells treated with CDDP and AMXT incrementally enhanced cell viability. More, when all three amino acids were supplemented, viability increased by about 20% (Fig. 5D), similar in the magnitude to the rescue observed with application of the ROS scavenger NAC.

**Figure 5.**
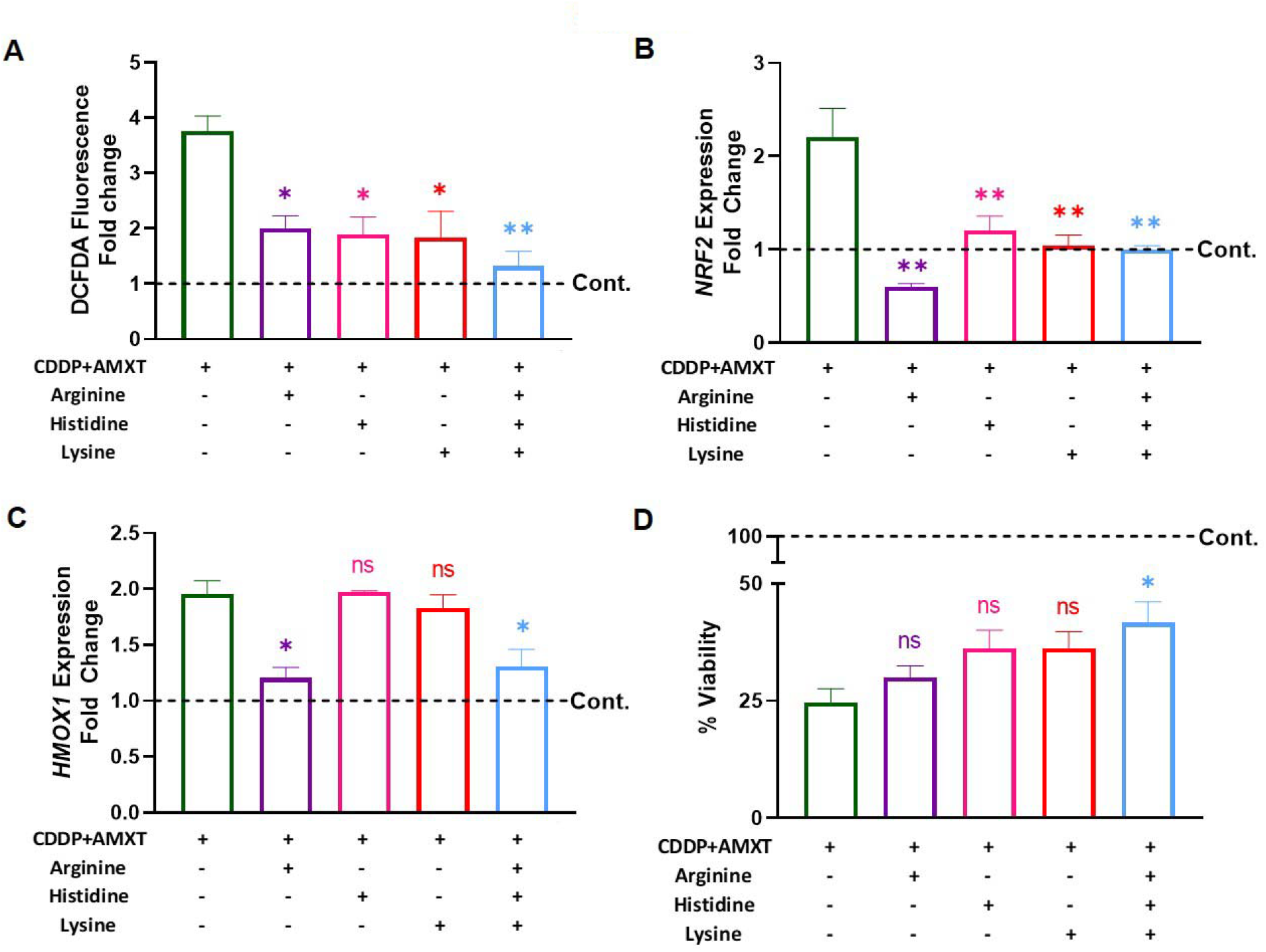
Amino acid supplementation reduces cell death and reactive oxygen species induced by cisplatin AMXT-1501 combination. (A) DCFDA fluorescence measured in MOC2 cells after treatment with 2 µM CDDP and 20 µM AMXT in the presence of arginine, histidine, and lysine for 24h. Expression of (B) *NRF2* and (C) *HMOX1* in MOC2 cells after treatment with CDDP and AMXT in the presence of arginine, histidine, and lysine for 24h. (D) Cell proliferation assay in MOC2 cells after treatment with CDDP and AMXT in the presence of arginine, histidine, and lysine for 72h. Comparisons are made between CDDP + AMXT treatment and other treatment groups. Statistical significance was calculated using one-way ANOVA with Tukey’s multiple comparison test. *p < 0.05, **p < 0.001.

## Discussion

Aggressive and recurrent head and neck cancer is driven by oxidative phosphorylation and ROS (35). As oxaliplatin has been shown to increase polyamine catabolism, we hypothesized that the combination of cisplatin and AMXT-1501, a polyamine transport inhibitor, would act synergistically to diminish tumor cell viability and proliferation. We observed both increased ROS and apoptotic cell death in cells treated with AMXT-1501 and cisplatin.

Interestingly, we found that the combination of AMXT-1501 and cisplatin did not result in altered intracellular polyamine levels. The combination of AMXT-1501 with standard-of-care cisplatin in head and neck cancer may offer a potent, viable option for treating aggressive head and neck cancers.

Polyamine blockade therapy (PBT) using AMXT-1501, a polyamine transport inhibitor, and difluoromethylornithine (DFMO), an ornithine decarboxylase inhibitor (39, 40), has shown promising results in preclinical models of diffuse intrinsic pontine glioma (2) and *MYCN* transgenic mouse models (40). In this study, we sought to combine a clinically implemented, standard-of-care agent for the treatment of head and neck cancer, cisplatin, with polyamine transport blockade, to examine whether the combination of cisplatin and AMXT-1501 could work harmoniously to deplete intracellular polyamines and abrogate tumor growth. Our data illustrated evidence of synergy between AMXT-1501 and cisplatin both *in vitro* and *in vivo* in head and neck cancer cells which was driven by ROS, but independent of intracellular polyamine levels.

Another important role of polyamines is redox regulation (41). Prior work found that chemically induced oxidative stress using tert-butylhydroquinone in human hepatoma Huh7 cells led to upregulation of ornithine decarboxylase and spermidine/spermine-N^1^-acetyltransferase, an effect mediated by NRF2, a transcription factor that regulates antioxidant response element (ARE) gene expression (42–44). On the other hand, *in vitro* work in human ovarian carcinoma A2780 cells exposed to cisplatin or oxaliplatin observed downregulation of *ODC1* and *SRM* with upregulation of *SSAT1* expression (24). Spermidine is known to serve as a substrate for hypusination of eukaryotic initiation factor 5A (eIF5A) through the rate-limiting enzyme, deoxyhypusine synthase. Hypusinated eIF5A then goes on to increase expression of mitochondrial proteins involved in oxidative phosphorylation (OXPHOS) (45). While we did not observe evidence of spermidine depletion in any of the CDDP, AMXT, or combination treatment conditions, oxidative stress was increased to a greater extent in the combination treatment relative to either treatment alone or control. Moreover, adding back spermidine to cell culture did not rescue cell death in the context of CDDP and AMXT combination therapy. Collectively, these data suggest a mechanism of cytotoxicity independent of spermidine levels or polyamine catabolism-related oxidative stress.

It has been observed that cationic amino acids, specifically arginine, reduce oxidative stress by stimulating cellular glutathione synthesis and activating NRF2 to upregulate ARE gene expression. Additionally, arginine acts as a superoxide scavenger (46, 47). Arginine is also known to be a precursor to ornithine and other downstream polyamines (48). Because of this, we hypothesized that arginine and other cationic amino acids may play a role in redox regulation in the setting of oxidative stress induced by the cisplatin and AMXT-1501 combination. We found that the presence of arginine, histidine, and lysine was able to significantly reduce ROS in our treated cells and partially recover cellular viability.

An advantage of our study is that the administration of cisplatin is routine in HNSCC patients, and AMXT-1501 is currently being tested in clinical trials. Administration of AMXT-1501 is relatively simple through oral formulation so combination of the two drugs is realistically feasible in the future. Limitations of our current study include the role of other genomic, epigenomic, and/or metabolic mechanisms improving tumor control with AMXT-1501 and cisplatin combination therapy remain unknown. This was outside the scope of this study, but potential mechanisms including differential competition for metabolites and the effect of combination AMXT-1501 and cisplatin on anti-tumor immunity remains to be clarified. Ongoing work aims to define the effect of AMXT-1501 and cisplatin on anti-tumor immunity.

## Methods

### Cell lines and reagents

MOC2, TAb2, and MEER PD-1 resistant (PD1R) cell lines were cultured in media as previously described and incubated at 37°C with 5% CO2 (25–27). All cell lines were authenticated and validated within 6 months of use. Cisplatin (CDDP) was purchased from EMD Millipore. AMXT-1501 (AMXT) and difluoromethylornithine (DFMO) were provided by Aminex Therapeutics. N-Acetyl Cysteine (NAC), MitoTEMPO (MT), spermidine, aminoguanidine, arginine, histidine, and lysine were purchased from Sigma. Elesclomol (ES) was purchased from Selleckchem.

### Cell proliferation assays

Cells were plated in 96-well plates at 1 – 2 x 10^3^ cells per well and were allowed to adhere overnight. Cells were then treated with the indicated doses of CDDP and AMXT. After 72 hours of treatment, 10 µL of premix WST-1 cell proliferation assay (Takara Bio, Inc.) was added to each well and the plate was incubated for an additional 2 hours. Absorbance was measured at 450 nm using a microplate reader (BioTek Instruments, Inc.). IC50 values were determined by a variable slope dose-response curve using GraphPad Prism software. To assess synergy, cells were treated with the combination of CDDP and AMXT at a ratio of 1:10 and CompuSyn software (https://www.combosyn.com) was used to calculate the Combination Indices (CI), where CI less than 1.0 indicates synergy, CI between 1.0 and 2.0 indicates additivity, and CI greater than 2.0 indicates antagonism. Where indicated, 3 mM of DFMO, 20 mM of NAC, 20 µM of MT, and 10 nM of ES were used. Additionally, 2300 µM of arginine, 850 µM of histidine, and 2300 µM of lysine were used, approximately 4-times the concentration of each in standard MOC media (49). When spermidine was used, 1 mM of aminoguanidine was also added to inhibit serum amine oxidase activity.

### Animal studies

Female C57BL/6 mice (The Jackson Laboratory) were injected subcutaneously with MOC2 (5 x 10^5^) or TAb2 (1 x 10^6^) cells in 30% Matrigel (Corning) in bilateral flanks. Mice were randomized after 7-10 days into control, CDDP, AMXT, and combination groups with 5-10 mice per group. Cisplatin was dosed at 3 mg/kg twice per week and administered intraperitoneally. AMXT-1501 dicaprate was dissolved in 3.3% mannitol and administered subcutaneously at 2.5, 5, or 10 mg/kg/day. The volumes of tumors were measured each day. Animals were handled and euthanized according to IACUC protocol. Tumors and kidneys were harvested, embedded in formalin, and given to the core facility for immunohistochemistry at the Pitt Biospecimen Core for H&E and TUNEL staining. Images were scanned at 20X magnification. H&E-stained kidney slides were examined and graded by a board-certified pathologist at HistoWiz (https://home.histowiz.com).

### Annexin V/PI flow cytometry

MOC2 cells were plated in a 6-well plate at 2.5 x 10^5^ cells per well and allowed to adhere overnight. Cells were then treated with either 2 µM CDDP, 20 µM AMXT, or combination. After 72 hours of treatment, cells were collected and prepared according to FITC Annexin V Apoptosis Detection Kit (BD Biosciences) protocol. Briefly, 1 x 10^5^ cells were washed once in PBS and resuspended in 100 µL Annexin V binding buffer. Cells were incubated in 5 µL FITC Annexin V and 5 µL PI for 15 minutes and then diluted to the appropriate volume of Annexin V binding buffer. Cells were analyzed using Beckman Coulter CytoFLEX.

### Measurement of reactive oxygen species

MOC2 cells were plated in a 96-well black well clear bottom plate at 8,000 cells per well and allowed to adhere overnight. Cells were then treated with either 2 µM CDDP, 10 nM ES, or 20 µM AMXT in the absence or presence of NAC, MT, or amino acids. After 24 hours of treatment, 2’,7’-dichlorodihydrofluorescein diacetate (DCFDA/H2DCFDA) Cellular ROS Assay (abcam) and MitoSOX Red Mitochondrial Superoxide Indicator (invitrogen) kits were used to measure relative levels of ROS in each treatment group according to the manufacturer’s protocol. Briefly, cells were incubated in 20 µM DCFDA or 5 µM MitoSOX solution. Fluorescence was measured (Magellan Pro Tecan) at 495/529 nm and background subtraction was performed at 485/535 nm for DCFDA assay and 510/580 nm for MitoSOX.

### Real-Time qPCR

MOC2 cells were plated in 6-cm dishes and allowed to reach 60% confluency. Cells were then treated with CDDP and AMXT in the absence or presence of NAC or amino acids. After 24 hours of treatment, cells were collected, and RNA was isolated using Rneasy kit according to manufacturer’s protocol. First-strand cDNA was made using iScript RT and qPCR was done with a suitable dilution of cDNA. Primer sequences for target genes are listed in supplemental table 1. RT conditions used were: 15s denaturation/95°C, 30s annealing/60°C, and 30s extension/72°C for 40 cycles. Relative gene expressions were calculated using the 2^-ΔΔCq^ method (50).

### High-performance liquid chromatography

MOC2 cells treated for 24 hours were collected, frozen in dry ice and ethanol, and sent to Johns Hopkins Sidney Kimmel Comprehensive Cancer Center. Perchloric acid-extracted lysates were labeled with dansyl chloride for detection using HPLC as previously described (51). Polyamine concentrations were quantified relative to total cellular protein in the lysate, as determined by Bradford assay (52).

### Statistics

All experiments were performed independently at least 3 times, each in triplicate. Unless otherwise stated, all graphs represent combined averages of each individual experiment. Bars are plotted as means ± SEM and are expressed as % fold change of the control. Statistical analysis was performed using GraphPad Prism.

### Author contributions

R.A.H. and U.D. designed research; A.Y., N.W., and J.F. performed research; T.M.S., M.B., and R.A.C. contributed new reagents/analytic tools; A.Y., T.M.S., R.A.C., R.A.H., and U.D. analyzed data; A.Y. and R.A.H. wrote the paper.

## Acknowledgements

We thank Dr. Greg M. Delgoffe and Dr. Jing Hong Wang at the University of Pittsburgh for providing us with MEER PD1R and TAb2 cell lines, respectively.

## Financial Support

This study is supported by grants I01 BX-003456, R01 DE028343, R00CA207871 and the Mosites Fund for personalized medicine to U.D.; grants R01CA226765, P41EB028239, R01CA204345, R01CA235863, Samuel Waxman Cancer Research Foundation, University of Pennsylvania Orphan Disease Center Million Dollar Bike Ride (MDBR-20-135-SRS), Chan Zuckerberg Initiative, and a research contract with Panbela Therapeutics Inc. to R.A.C. and T.M.S.; American Head and Neck Society / American Academy of Otolaryngology – Head & Neck Surgery Foundation Young Investigator Combined Award to R.A.H. This publication was also made possible by the Johns Hopkins Institute for Clinical and Translational Research (ICTR) which is funded in part by grant KL2TR003099 from the National Center for Advancing Translational Sciences (NCATS) a component of the National Institutes of Health (NIH), and NIH Roadmap for Medical Research. Its contents are solely the responsibility of the authors and do not necessarily represent the official view of the Johns Hopkins ICTR, NCATS or NIH.

**Supplemental Figure 1.**

(A) Cell proliferation measured every 24 hours in untreated MOC2 cells. Measurements are normalized to proliferation at time 0-hours and plotted as relative fold change.

**Supplemental Figure 2.**

(A) Cell proliferation assay in MOC2 cells after treatment with 3 mM DFMO, 2 µM CDDP, and 20 µM AMXT for 72h in media supplemented with 1 µM spermidine and 1 mM aminoguanidine. Comparisons made are indicated by lines connected to bars. Statistical significance was calculated using one-way ANOVA with Tukey’s multiple comparison test.

**Supplemental Figure 3.**

H&E stain of harvested kidneys of mice after treatment with (A) 10 mg/kg/day AMXT and (B) CDDP + AMXT displaying areas of tubular basophilia (white arrows) and individual necrotic tubular cells (black arrows).

**Supplemental Figure 4.**

(A) Tumor growth curves in mice xenografted with MOC2 cells after treatment with CDDP and 10 mg/kg/day of AMXT. Treatment was discontinued after day 11. (B) Images for harvested MOC2 tumors are shown for each treatment group. For A, comparisons are made between CDDP + AMXT treatment and individual treatment groups. Statistical significance was calculated using two-way ANOVA with Tukey’s multiple comparison test. ***p < 0.0001.

**Supplemental Figure 5.**

(A) Bar graph analysis of TUNEL staining in harvested TAb2 tumors after treatment with CDDP and AMXT. Bars display the average number of TUNEL-positive cells per high-powered field. (B) Representative TUNEL stain images are shown. For A, comparisons are made between CDDP + AMXT treatment and individual treatment groups. Statistical significance was calculated using one-way ANOVA with Tukey’s multiple comparison test. *p < 0.05, ***p < 0.0001.

**Supplemental Figure 6.**

(A) DCFDA fluorescence in MOC2 cells after treatment with 10 nM ES and 20 µM for 24h. (B) Cell proliferation assay in MOC2 cells treated with 10 nM ES and 20 µM AMXT for 72h. Comparisons are made between ES + AMXT treatment and individual treatment groups. Statistical significance was calculated using one-way ANOVA with Tukey’s multiple comparison test. *p < 0.05, **p < 0.001.

**Supplemental Figure 7.**

(A) Cell proliferation assay after treatment with CDDP and AMXT in media supplemented with indicated concentrations of spermidine and 1mM aminoguanidine. Intracellular levels of (B) putrescine, (C) spermidine, and (D) spermine after treatment with CDDP and AMXT for 24h.

**Supplemental Table 1.**

List of qPCR primers.

