## Supplemental Figures for "Polyamine transport inhibition and cisplatin synergistically enhance tumor control through oxidative stress in murine head and neck cancer models"

### Supplemental Figure 1

A

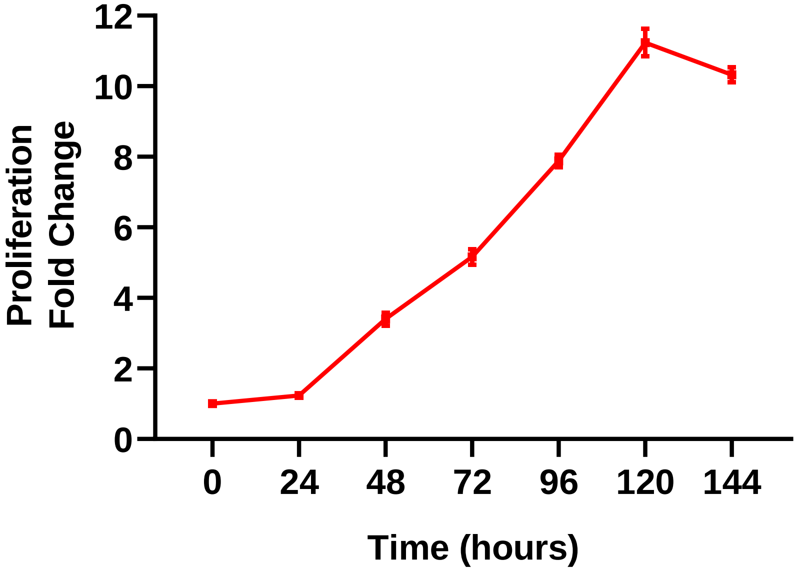

### Supplemental Figure 2

A

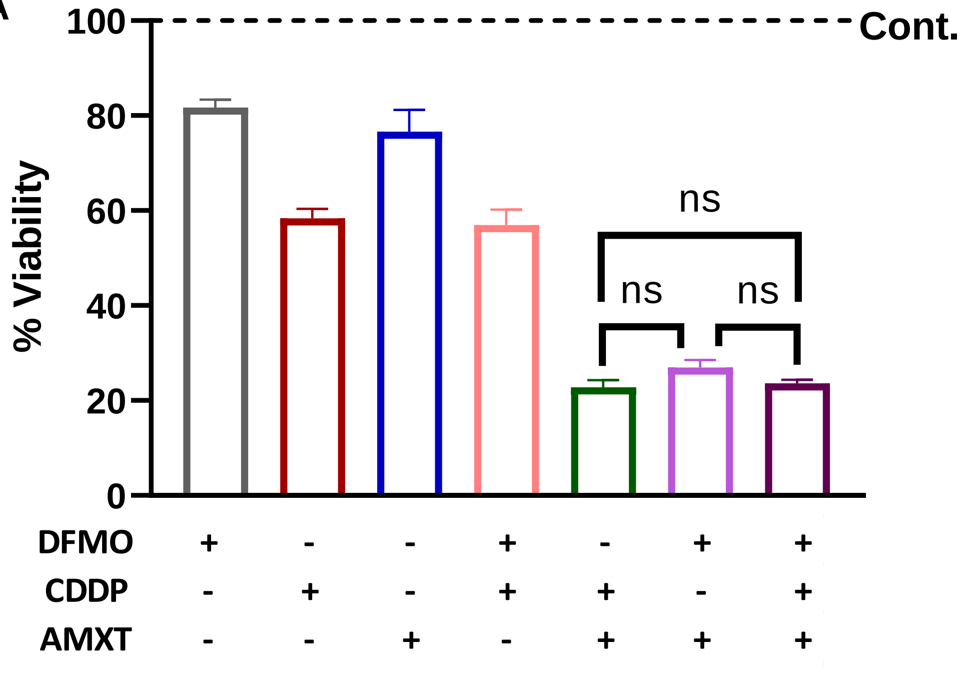

### Supplemental Figure 3

**A**

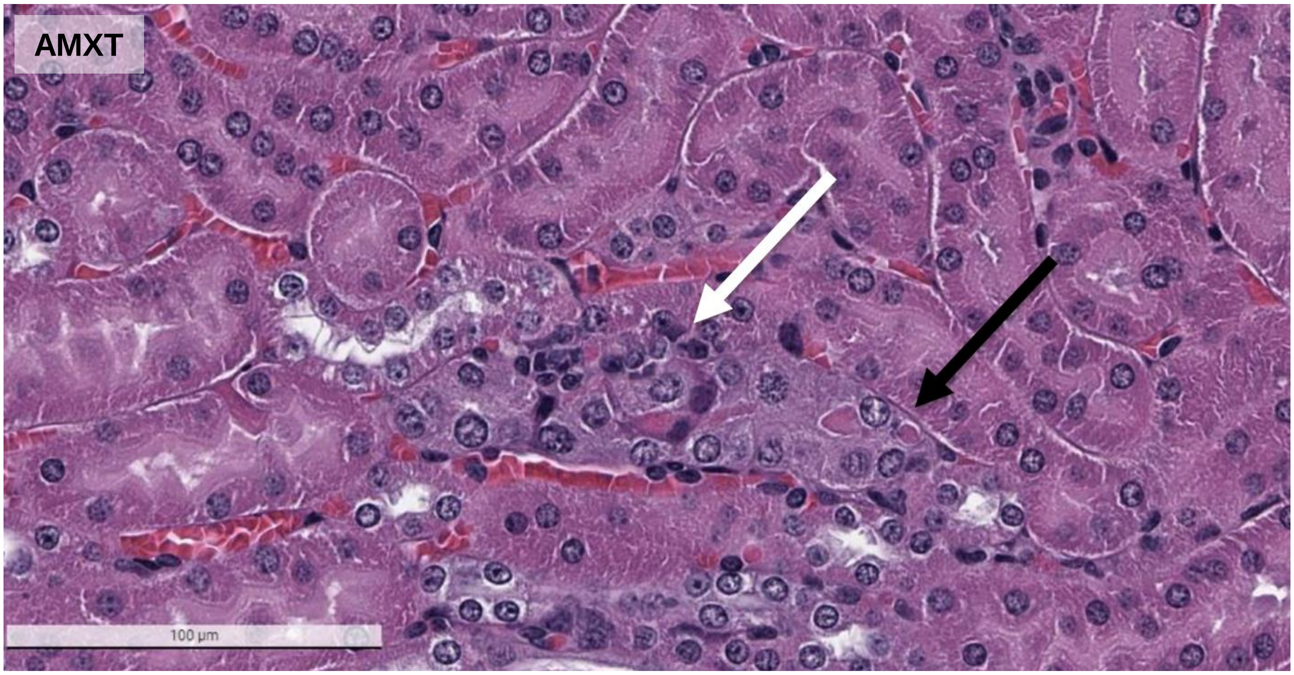


**B**

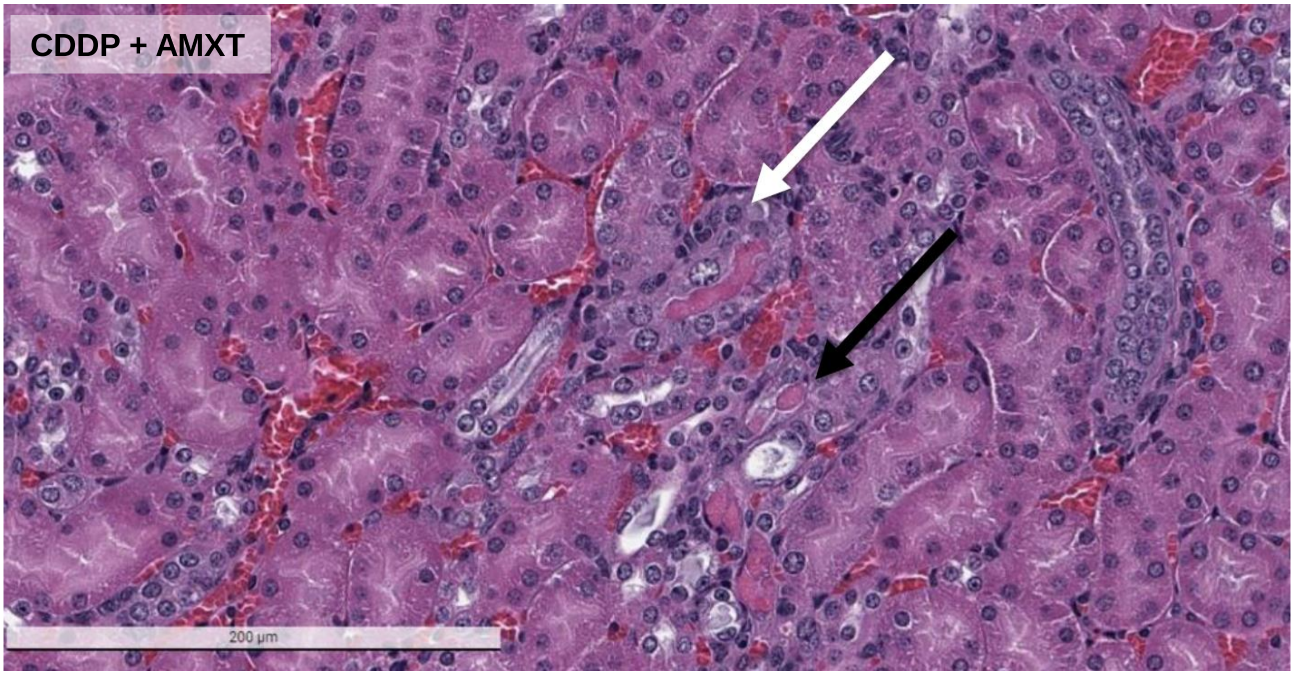

#### Supplemental Figure 4

A

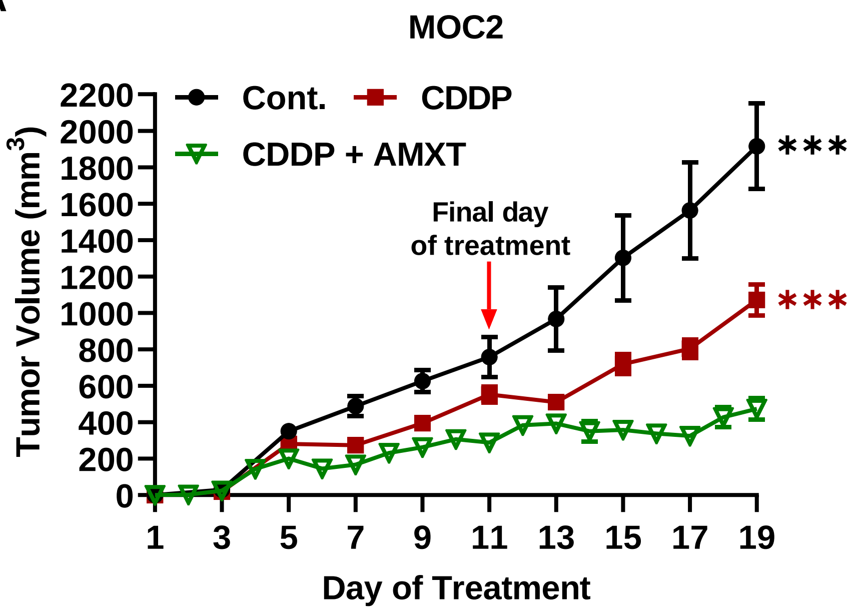


B

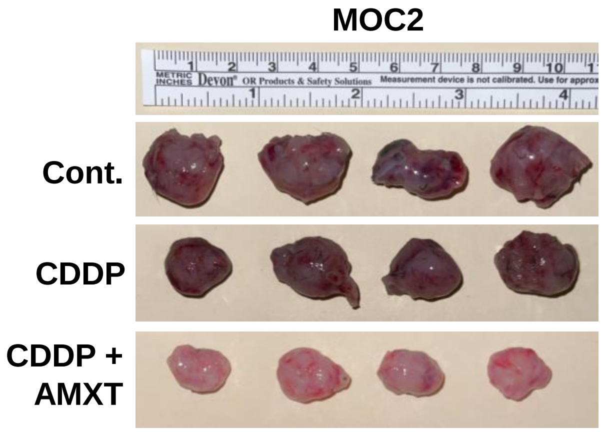

### Supplemental Figure 5

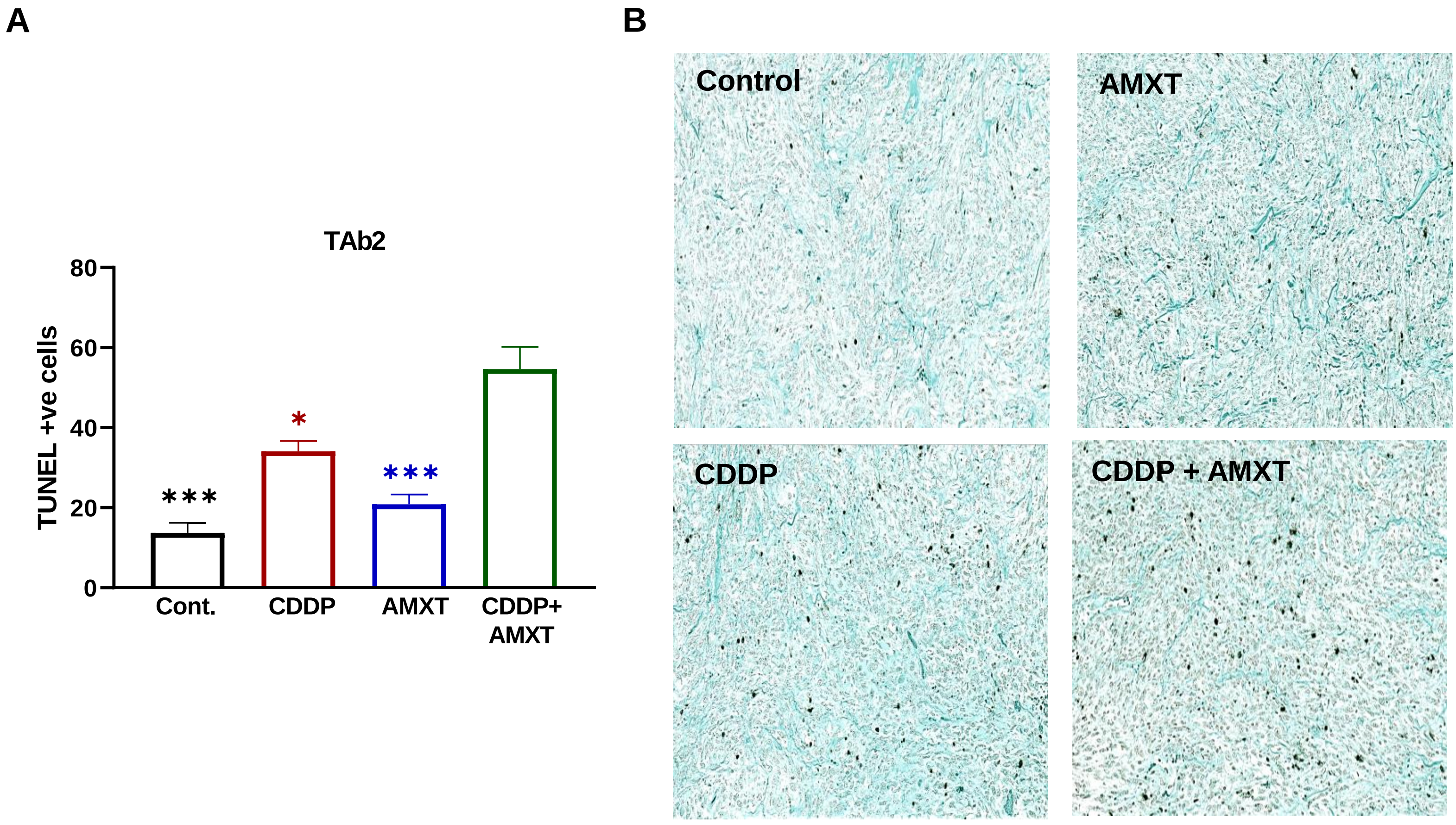

### Supplemental Figure 6

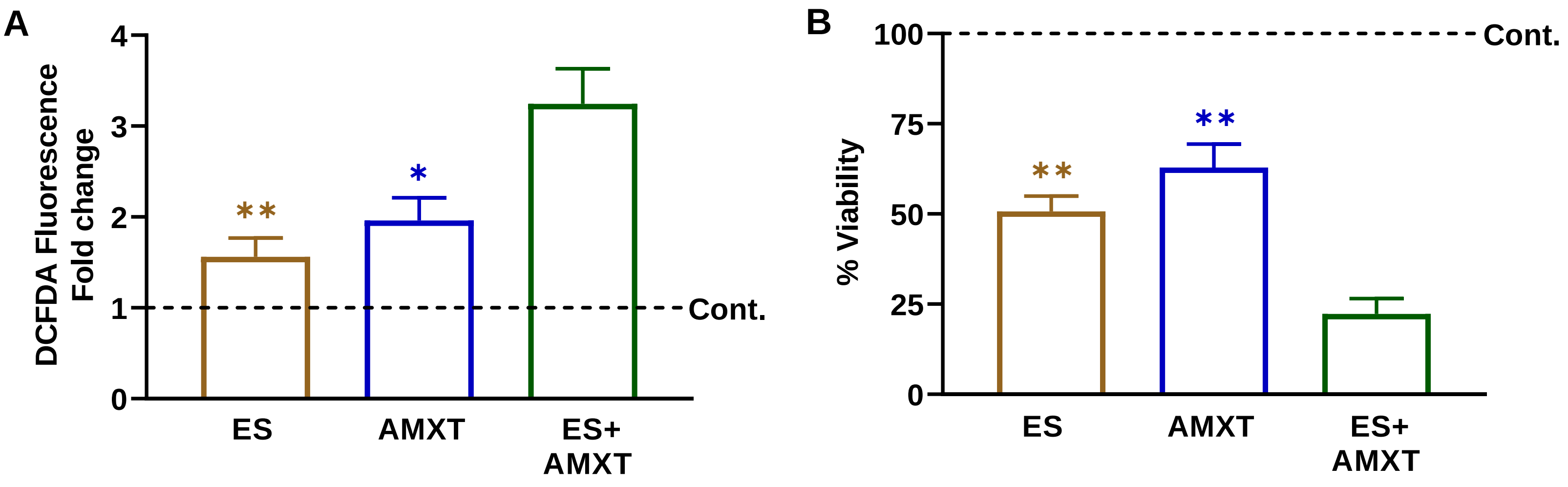

#### Supplemental Figure 7

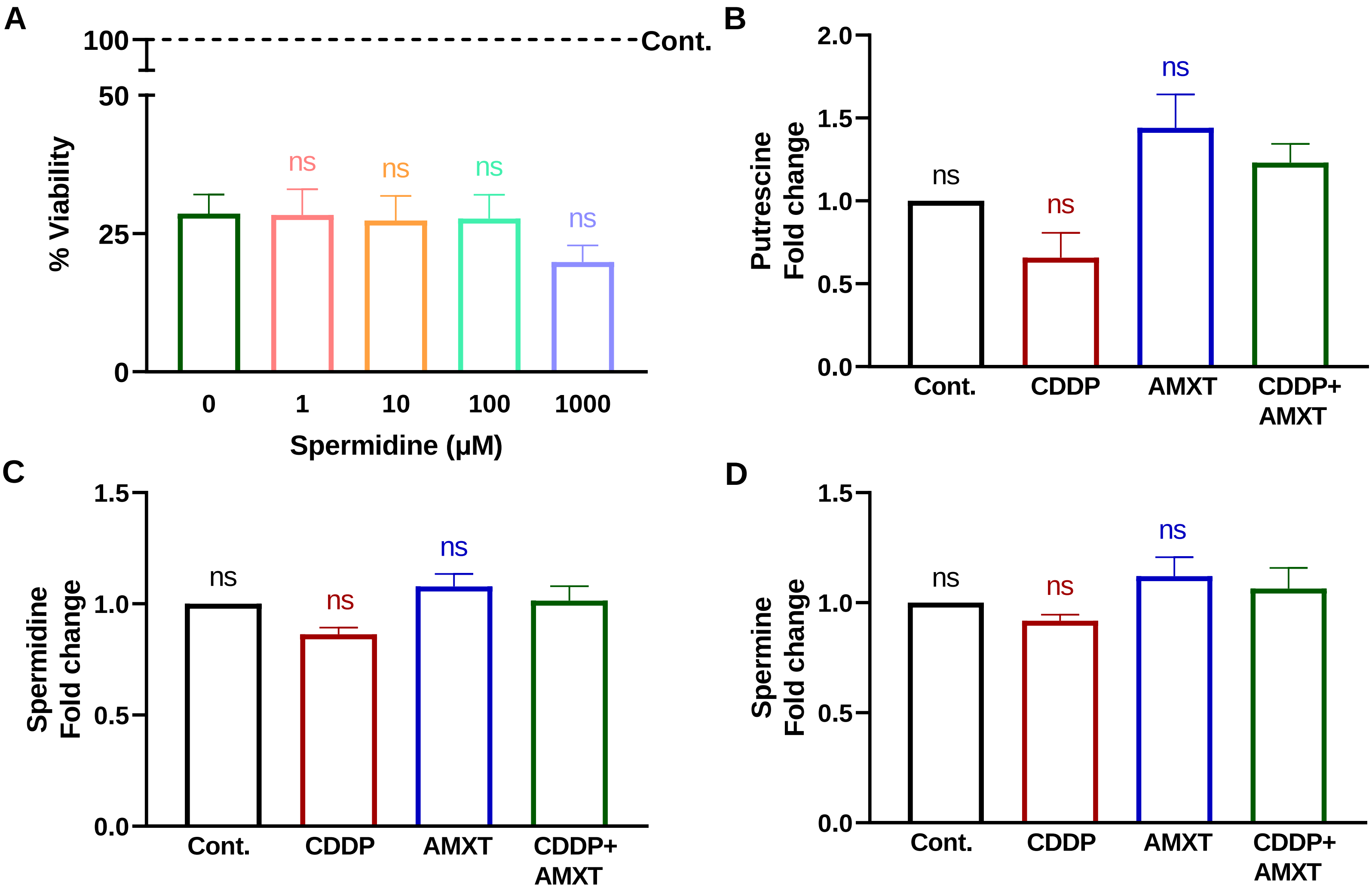

### Supplemental Table 1

| Gene | | Sequence |
|---|---|---|
| *NRF2* | Forward | 5’-CAAGACTTGGGCCACTAAAAAGAC-3’ |
| | Reverse | 5’-AGTAAGGCTTTCCATCCTCATCAC-3’ |
| *HMOX1* | Forward | 5’-CACTTCGTCAGAGGCCTGCTA-3’ |
| | Reverse | 5’-GTCTGGGATGAGCTAGTGCTGAT-3’ |
| *GAPDH* | Forward | 5’-AGGTCGGTGTGAACGGATTTG-3’ |
| | Reverse | 5’-GGGGTCGTTGATGGCAACA-3’ |
| *Actin* | Forward | 5’-CGGTTCCGATGCCCTGAGGCTCTT-3’ |
| | Reverse | 5’-CGTCACACTTCATGATGGAATTGA-3’ |
